# New Insight Into the Neuroimmune Interplay In *Pseudomonas aeruginosa* Keratitis

**DOI:** 10.1101/2025.03.06.641908

**Authors:** Naman Gupta, Giovanni LoGrasso, Linda D. Hazlett, Shunbin Xu

**Author notes:** Correspondence: Shunbin Xu, MD/PhD. Department of Ophthalmology, Visual and Anatomical Sciences, Wayne State University School of Medicine/Kresge Eye Institute, Detroit, MI48203.

## Abstract

**Purpose:** The miR-183/96/182 cluster (miR-183C) is required for normal functions of sensory neurons (SN) and various immune cells, including myeloid cells (MC). This research aims to reveal the roles of miR-183C of SN in the interplay of corneal sensory nerves (CSN) and MC during *Pseudomonas aeruginosa* (PA) keratitis.

**Methods:** Double-tracing mice with SN-specific (SNS) conditional knockout of miR-183C (CKO) and age-and sex-matched wild type (WT) controls were used. Their CSN are labeled with Red Fluorescent Protein (RFP); MC with Enhanced Green (EG)FP. The left corneas were scarified and infected with ATCC19660 PA. Corneal flatmounts were prepared at 3, 6, and 12 hours post-infection (hpi) and 1, 3, and 5 days (d)pi for confocal microscopy. Myeloperoxidase (MPO) assay and plate count were performed at 1 dpi.

**Results:** In WT mice, CSN began to degenerate as early as 3 hpi, starting from the fine terminal CSN in the epithelial/subepithelial layers, most prominently in the central region. By 1 dpi, CSN in the epithelium/subepithelial layer were nearly completely destroyed, while stromal nerves persisted. From 3 dpi, CSN were obliterated in both layers. In CKO vs WT mice, CNS followed a slightly slower pace of degeneration. CSN density was decreased at 3 and 6 hpi. However, at 3 dpi, residual large-diameter stromal CSN were better preserved. MC were decreased in the central cornea at 3 and 6 hpi, but increased in the periphery. Both changes were more prominent in CKO vs WT mice. At 12 hpi, densely packed MC formed a ring-shaped band circling a “dark” zone nearly devoid of MC, colocalizing with CSN most degenerated zone in the central cornea. In CKO vs WT, the ring center was larger with fewer MC. At 1 dpi, the entire cornea was filled with MC; however, MC density was lower in CKO mice. An MPO assay showed decreased neutrophils in PA-infected cornea of CKO mice. This led to a decreased severity of PA keratitis at 3 dpi.

**Conclusions:** This double-tracing model reveals the interplay between CSN and MC during PA keratitis with greater clarity, providing new insights into PA keratitis. CSN-imposed modulation on innate immunity is most impressive within 24 hours after infection. Functionally, the miR-183C in CSN modulates CSN density and the dynamics of MC fluxes-a neuroimmune interaction in display.

## Introduction

*Pseudomonas aeruginosa* (PA) is a Gram-negative bacterium and a common causative organism associated with contact lens-related disease in developed countries^1^. PA-induced keratitis is one of the most rapidly developing and destructive diseases of the cornea^1^. It is estimated that 140 million people wear contact lenses worldwide, making PA keratitis a global cause of visual impairment and blindness^1–5^. Currently, PA keratitis is mainly treated by topical administration of antibiotics; in severe cases, subconjunctival injection may be employed. Although antibiotic treatment reduces the bacterial burden, tissue damage often occurs as a result of a poorly controlled host immune response^6, 7^. Additionally, frequent emergence of antibiotic resistant strains poses serious challenges for the effective management of the disease^8–10^. Deeper insights into the pathogenesis will be the keys to the development of alternative treatment of the disease.

The cornea is the most densely sensory innerved tissue in the body^11–13^. The neuronal cell bodies of corneal sensory nerves (CSN) reside in the trigeminal ganglion (TG) and can be activated by PA^14^. Evidences suggest that CSN secrete neuropeptides, e.g., substance (s) P and Calcitonin Gene-Related Peptide (CGRP), which modulate the immune response to PA infection^14–19^. Although a few studies indicated that CSN were quickly degenerated upon PA infection^14, 20^, the data in these reports were limited and fragmented and lacked details of the dynamic changes of CSN. The simultaneous changes of the CSN and innate immune cells have not been studied hindering deeper understanding of the mechanisms of neuroimmune interaction during PA keratitis.

microRNAs (miRNAs) are small, endogenous, non-coding RNAs and are important post-transcriptional regulators of gene expression^21–24^. They play an important role in human diseases^25–32^ and are viable therapeutic targets^33–36^. However, their role in bacterial keratitis remain largely unexplored. Previously, we identified a conserved, paralogous miRNA cluster, the miR-183/96/182 cluster (referred to as miR-183C from here on), which is highly expressed in all primary sensory neurons of all major sensory domains, including the trigeminal ganglia (TG), where sensory neurons innervating the cornea reside^37–39^. Inactivation of the miR-183C disrupts the highly organized whorl-like pattern of the subbasal plexus of the CSN and results in decreased CSN density, sensitivity to mechanical stimuli and reduced levels of neuropeptides in the cornea^39, 40^. Furthermore, we discovered that the miR-183C is also expressed in innate myeloid cells (MC), including corneal resident myeloid cells (CRMC)^41^ as well as circulating MC, e.g., macrophage (Mφ) and polymorphonuclear leukocytes (PMN)^39^. miR-183C regulates the production of pro-inflammatory cytokines in response to PA infection by these cells and their phagocytosis and bacterial killing capacity through targeting key genes involved in related pathways^39–42^. Complete inactivation of miR-183C in a conventional knockout mouse model results in decreased immune/inflammatory response and reduced severity of PA keratitis^39^. Topical application of anti-miR-183C in the cornea leads to a reversible CSN regression, enhanced functional maturation of infiltrating MC and a decreased severity of PA keratitis^43^, suggesting that miR-183C is a therapeutic target for the treatment of PA keratitis. Recently, we developed a sensory neuron-specific (SNS) and a myeloid cell-specific (MS) conditional miR-183C knockout mouse models^40^. Using these models, we demonstrated that, in naïve mice, miR-183C imposes both intrinsic and extrinsic regulations on CSN, CRMC and corneal function through cell type-specific target genes^40^. These data support the hypothesis that miR-183C modulates corneal responses to PA infection through its dual regulation of CSN and innate immune cells and neuroimmune interaction. In the SNS-CKO mouse model, CSN are clearly labeled with Nav1.8-Cre-induced SNS expression of TdTomato red fluorescent protein (RFP), while the MC with Csf1r-promoter-driven enhanced green fluorescent protein (EGFP)^40^. This double-tracing mouse model provides an unprecedented powerful tool to simultaneously monitor CSN and MC, including both resident and infiltrating MC^40^. Here we report our observations of the simultaneous, dynamic changes of CSN and MC during PA keratitis and the impact of SNS-CKO of miR-183C. Our data provide new insights into the neuroimmune interaction and the pathogenesis of PA keratitis and the roles of miR-183C in CSN in the development of the disease.

## Methods

### Mice

All experiments and procedures involving mice and their care were pre-reviewed and approved by the Wayne State University Institutional Animal Care and Use Committee and carried out in accordance with National Institute of Health and Association for Research in Vision and Ophthalmology (ARVO) guidelines. The study is reported in accordance with ARRIVE guidelines. Euthanasia was performed by cervical dislocation under anesthesia with isoflurane followed by thoracotomy.

Mice with miR-183C CKO allele, the miR-183C^f/f^, were provided by Dr. Patrick Ernfors, Karolinska Institutet, Sweden through the European Mouse Mutant Archive (EMMA. ID: EM12387). The miR183C^f^ allele has two loxP sites flanking the 5’ and 3’ ends of the miR-183C for robust Cre-mediated miR-183C knockout^44^. The sensory nerve-specific Nav1.8-Cre mice^45^ were kindly provided by Dr. John N. Wood, University College London, through Dr. Theodore J. Price, University of Texas, Dallas. This strain is now available at the Jackson Laboratory (Stock Number: 036564). The voltage-gated sodium channel Na_v_1.8 (encoded by the Scn10a gene) is one of the signature genes of the majority of nociceptive sensory neurons in the TG and dorsal root ganglia (DRG)^14, 46–48^. Na_v_1.8 promoter-driven Cre recombinase (Na_v_1.8-Cre) is expressed in nearly all corneal nociceptive sensory nerves^14, 45, 49^.

### The Csf1r-EGFP or MacGreen mice^50^ were purchased from the Jackson lab (Stock number. 018549)

In this strain, the EGFP transgene is under the control of the 7.2-kb mouse colony stimulating factor 1 receptor (Csf1r) promoter, allowing specific expression of EGFP in the mononuclear phagocyte system (MPS) myeloid cells, including monocytes, Mj and dendritic cells (DCs)^50, 51^. The reporter strain, R26^LSL-RFP(+/+)^ mice^52^, also known as Ai14 (Stock number: 007914, the Jackson Laboratory) has a loxP-flanked STOP cassette (LSL) in front of a tdTomato RFP cassette, all of which are inserted into the ROSA26 locus^52^. The LSL prevents the transcription of tdTomato RFP; however, when Cre recombinase is present, the LSL cassette will be excised to allow the expression of tdTomato RFP. SNS-CKO mice [Na_v_1.8-Cre(+/-);miR-183C^f/f^;R26^LSL-RFP(+/+)^;Csf1r-EGFP(+/+)] and their WT littermates [Na_v_1.8-Cre(+/-);miR-183C^+/+^;R26^LSL-RFP(+/+)^;Csf1r-EGFP(+/+)] were produced by breeding of Na_v_1.8-Cre (+/-), miR-183C^f/f^, R26^LSL-RFP^ and Csf1r-EGF(+/-) mice. All breeding showed a normal mendelian inheritance pattern. 8-20 weeks old, male or female mice were used separately in all experiments. The age, sex and number of the mice in each experiment are specified in the figure legends and/or the text. On average, 5 mice/sex/genotype was used at each time-point for various analyses.

### PA infection

PA infection was conducted as we described previously^39, 41, 43^. Briefly, mice were anesthetized with isoflurane in a well-ventilated hood. The cornea of the left eye was scarified with a 25 5/8-gauge needle. Three 1-mm incisions were made to the corneal surface, which penetrated the epithelial layer, but no deeper than the superficial stroma. 5.0 × 10^6^ colony forming units (CFU) PA (strain 19660; ATCC) in a 5-μl volume was topically delivered. Corneal disease was graded at 1, 3 and 5 days-post-infection (dpi) using an established scale^15^: 0, clear or slight opacity, partially or fully covering the pupil; +1, slight opacity, covering the anterior segment; +2, dense opacity, partially or fully covering the pupil; +3, dense opacity, covering the entire anterior segment; and +4, corneal perforation. Photography with a slit lamp was used to illustrate disease at 1, 3 and 5 dpi. The corneas were harvested at 3, 6, and 12 hours post-infection (hpi) and 1, 3, and 5 days post-infection (dpi) for corneal flatmount preparation or other assays as detailed below.

### Corneal flatmount preparation

Corneal flatmount was performed as described previously^40, 41,43^. Briefly, mice were euthanized; eyes were enucleated and transferred to cold 1% paraformaldehyde (PFA) in 0.1 M phosphate buffer (PB), pH 7.4 for 10 minutes before being carefully dissected under a dissection microscope (VWR International, Radnor, PA). The cornea was carefully dissected out and fixed in cold 1% PFA in 0.1 M PB, pH 7.4 for another hour (h) on ice. The cornea was flattened by six evenly spaced cuts from the periphery toward the center and mounted in Vectashield media with DAPI (Vector Laboratories, Burlingame, CA) on Superfrost Plus slides (Fisherbrand).

### Quantification of corneal nerve density and myeloid cells

All slides were imaged using a TCS SP8 laser confocal microscope (Leica Microsystems Inc., Buffalo Grove, IL). To capture the images of the entire cornea, a series of Z-stacked images under 10x objective were taken across the entire cornea and stitched together. All images were taken and processed under the same parameters, e.g., laser power and digital gain and offset, without saturation to ensure that all images were comparable and the differences among samples reflected their biological differences. These stitched Z-stacked images were merged/flattened for cell counting or CSN density quantification using Adobe Photoshop CS6 (64 bit) and ImageJ 1.52p software (http://imagej.nih.gov/ij. NIH, Bethesda, MD, USA) as described previously^41^. Briefly, the raw image was first converted to the 16-bit grayscale; then the black & white binary image is optimized for the threshold to faithfully represent the original image and the cell density. The EGFP+ cells in the entire cornea were counted for corneas at 3 and 6 dpi. For corneas at 12 hpi and 1, 3, and 5 dpi, since the number of EGFP+ MC are too densely packed to be distinguished individually, we cannot reliably count the numbers; instead, we measured the volumes of fluorescence of comparable areas (Adobe Photoshop CS6) as relative quantification of MC for comparisons between WT and CKO samples.

To quantify CSN density, the mean value of pixels in a 500x500 μm^2^ square covering the whorl center of the subbasal plexus was quantified as the nerve density of the center. In the periphery, three 500x500 μm^2^ squares were randomly placed and measured; and the average was recorded as CSN density of the peripheral cornea.

### Myeloperoxidase (MPO) assay

A myeloperoxidase (MPO) assay was performed as described before to quantify the number of neutrophils^39, 43^. Briefly, corneas were harvested, homogenized in 1 ml potassium phosphate buffer (50 mM, pH 6.0) containing 0.5% hexadecyltrimethyl-ammonium bromide (Sigma-Aldrich). After four freeze/thaw cycles, 100 μl supernatant was added to 2.9 ml o-dianisidine dihydrochloride substrate buffer (16.7 mg/ml) with 0.0005% hydrogen peroxide. The change in absorbance at 460 nm was read every 30 seconds (s) for 5 min on a Helios-α spectrophotometer (ThermoFisher). Units of MPO per cornea were calculated: 1 unit of MPO activity is equivalent to ∼2 x 10^5^ PMN^53^.

### Quantitation of viable bacteria

Bacteria were quantitated by plate count assay as described before^39, 43^. Briefly, each cornea was homogenized in 1.0 ml sterile saline containing 0.25% BSA. 0.1 ml of the corneal homogenate was serially diluted 1:10 in the same solution and selected dilutions were plated in triplicate on *Pseudomonas* isolation agar (Becton Dickinson). Plates were incubated overnight (O/N) at 37 °C and the number of colonies counted. Results are reported as log_10_ number of CFU/cornea ± SEM.

**Statistical analyses** were performed as we described previously^40^. Briefly, When the comparison was made among more than 2 conditions, one-way ANOVA with Bonferroni’s multiple comparison test was employed (GraphPad Prism); adjusted p<0.05 was considered significant. Otherwise, a two-tailed Student’s t test was used to determine the significance; p<0.05 was considered significant. Each experiment was repeated at least once to ensure reproducibility and data from a representative experiment are shown. Quantitative data is expressed as the mean ± SEM.

## Results

### Dynamics of corneal sensory nerve degeneration during PA keratitis

To uncover the behaviors of CSN during PA infection, we infected (ATCC 19660) the left eyes of SNS-CKO and age-and sex-matched WT control mice and performed flatmount confocal imaging at 3, 6, and 12 hpi and 1, 3, and 5 dpi. Our data showed that CSN began to degenerate as early as 3 hpi, the earliest time-point of our study, in both WT and SNS-CKO mice (**Fig.1**. *: p<0.05 in **Fig.1E**). At 3 hpi, significant reduction of CSN density occurred first in the central region of the infected vs non-infected contralateral cornea; however, no significant change was observed in the peripheral cornea in both WT and SNS-CKO mice (**Fig.1A-E**).

**Figure 1.**
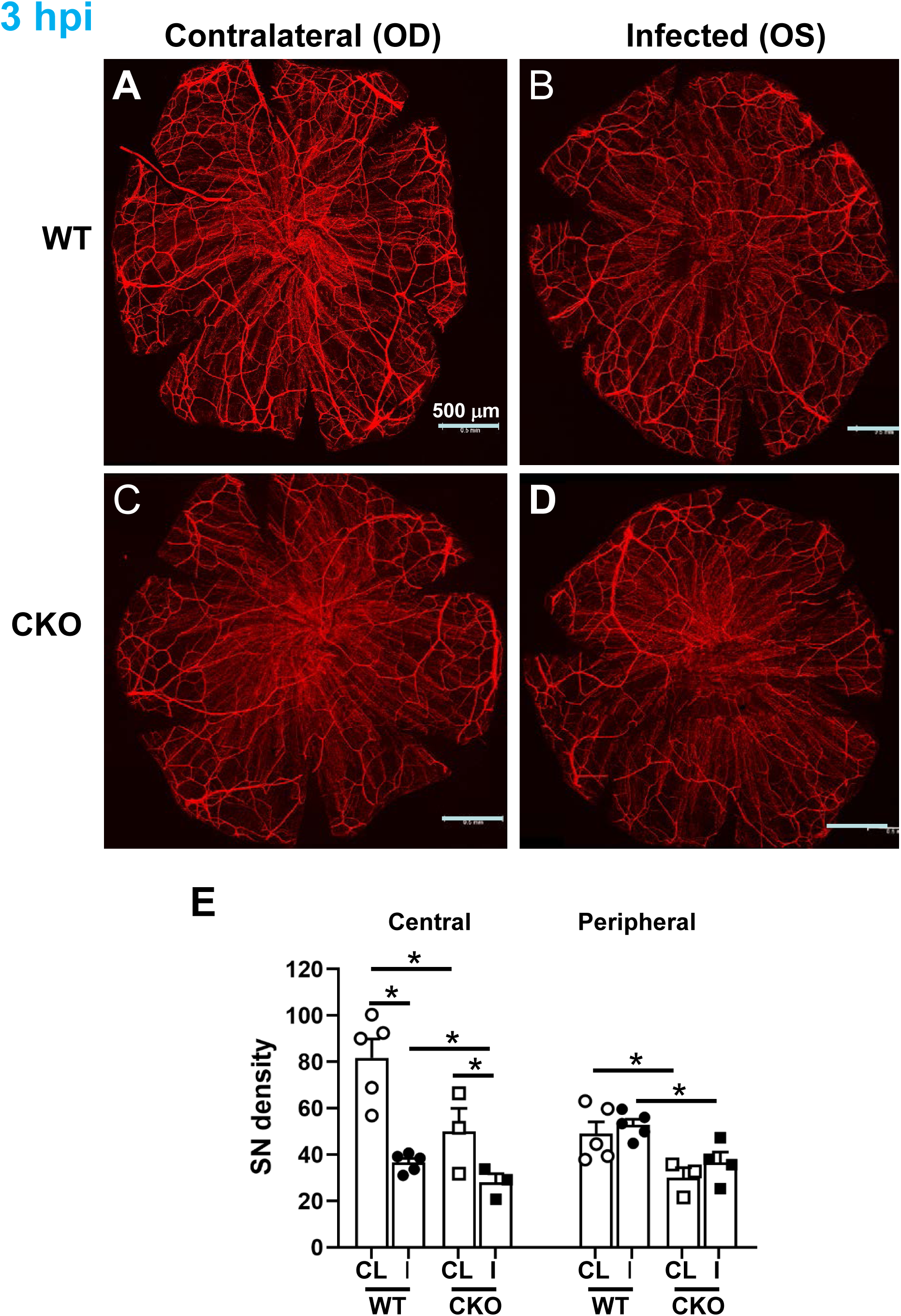
CSN began to degenerate as early as 3 hpi. **A-D**. Representative compressed flatmount confocal images of the corneas of PA infected (B,D) and non-infected contralateral eyes (A,C) of WT (A,B) and SNS-CKO mice (12-20 weeks old, female). OD: oculus dexter (right eye); OS: oculus sinister (left eye). Scale bars: 500 μm. **E**. Quantification of sensory nerve density of the central and peripheral regions of the cornea of WT and SNS-CKO mice. CL: contralateral eye; I: infected eye. *: p<0.05.

In WT mice, as the disease developed, the degeneration of CSN gradually worsened from the central region to the periphery and from the fine, terminal nerves in the epithelial/subepithelial layer to the larger stromal nerves (**Fig.2A**, blue line **Fig.3**). In the peripheral region, CSN degenerated in a slower pace when compared to the central region; significant degeneration did not occur until 12 hpi [**Fig.2A-c, Fig.3B** (^: p<0.05, 12 hpi vs 3 hpi)]. From 12 hpi to 1 dpi, CSN continued to degenerate in both central and peripheral regions [**Fig.2A**, **Fig.3** (‡: p<0.05 1 dpi vs 12 hpi)]. By 1 dpi, most of the fine terminal nerves in the epithelial and subepithelial layer were degenerated, while most of the stromal nerves were still preserved (**Fig.2A-d**). From 1 to 3 dpi, CSN degeneration in the peripheral region drastically accelerated in WT (blue line in **Fig.3B**. ∏: p<0.05, 3 dpi vs 1 dpi). From 3 dpi, all sensory nerves were nearly completely destroyed in both the central and peripheral regions of the WT mice (**Fig 2.Ae,f; Fig.3**). CSN in the contralateral eyes did not have significant changes at all times (data not shown).

**Figure 2.**
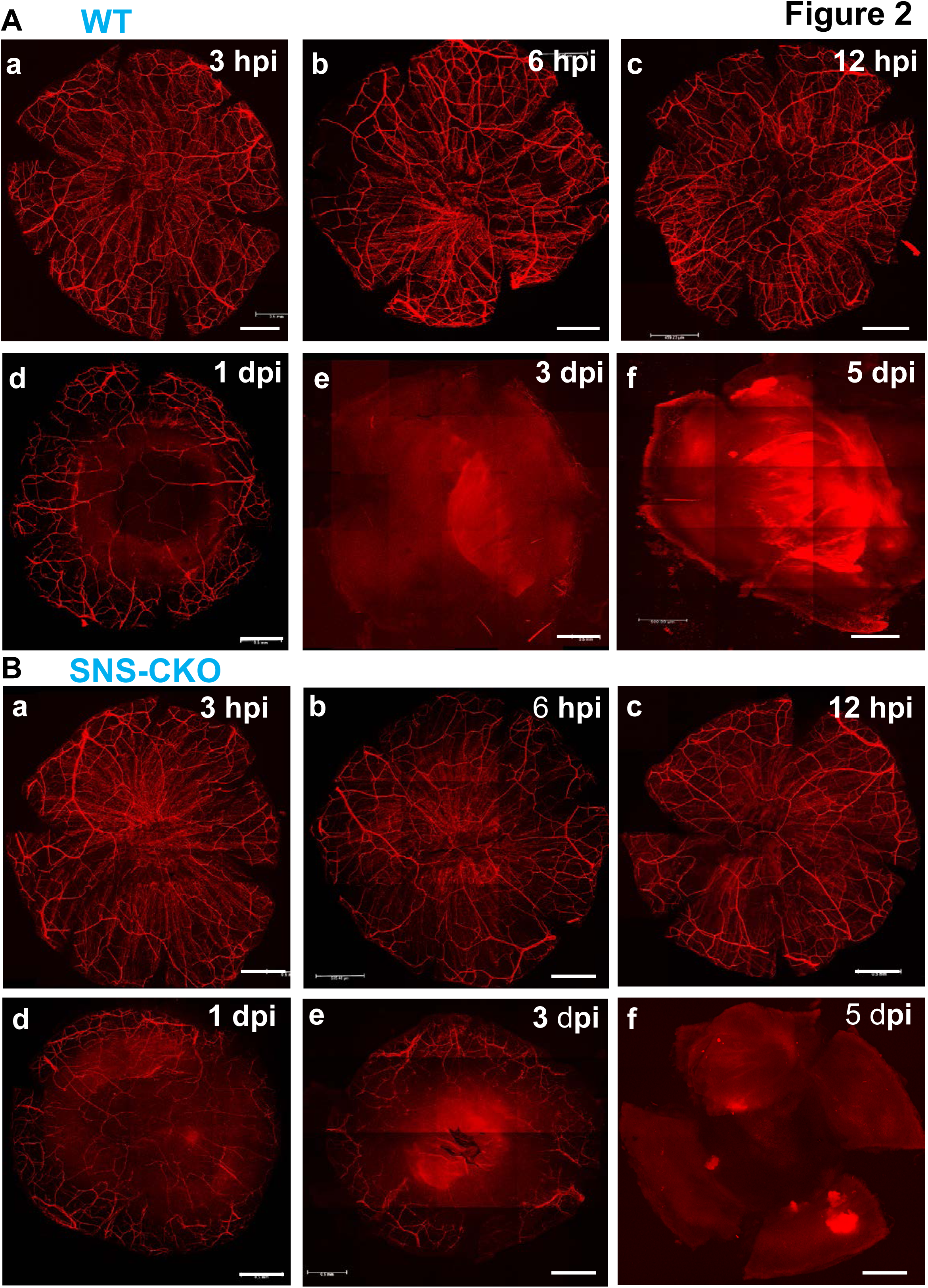
Representative compressed flatmount confocal images of the corneas of PA infected left eyes of WT (**A**) and SNS-CKO mice (**B**) at 3, 6 and 12 hpi and 1, 3 and 5 dpi. Scale bars: 500 μm.

**Figure 3.**
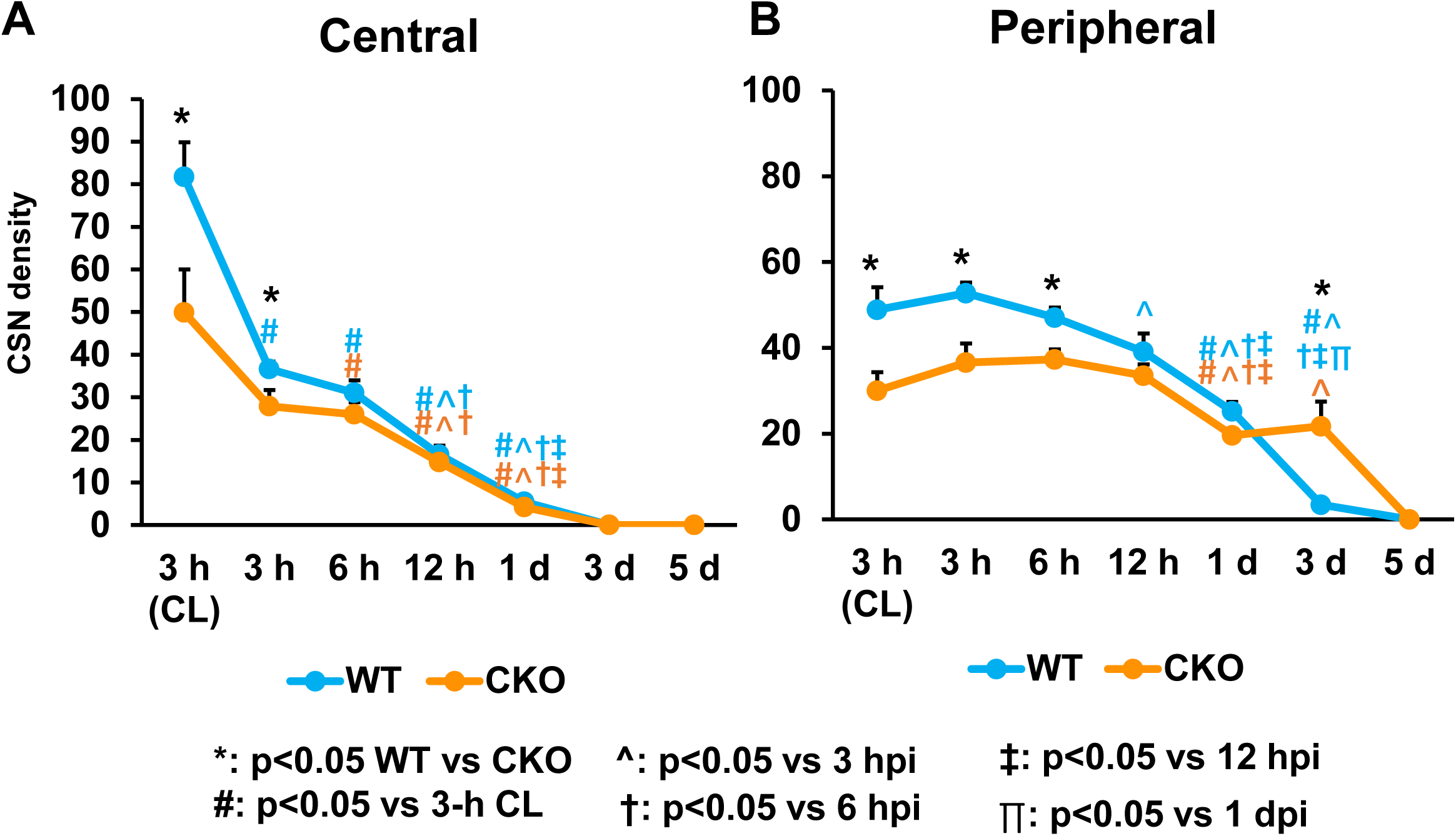
Dynamic changes of CSN density of the central (**A**) and peripheral regions of the corneas (**B**) of SNS-CKO and their age-and sex-matched WT controls at 3, 6, and 12 hours (h) and 1, 3, and 5 days post-infection (d), in comparison to the ones of the non-infected, contralateral eye at 3 hpi [3 h (CL)]. *: p<0.05, WT vs CKO; #: p<0.05, vs 3 h CL; ^: p<0.05, vs 3 hpi; †: p<0.05, vs 6 hpi; ‡: p<0.05, vs 12 hpi. ∏: p<0.05, vs 1 dpi. n= 5, 4, 6, 5, 5, and 5 for WT, and 3, 4, 5, 6, 5, and 5 for SNS-CKO at 3, 6, and 12 hpi, 1, 3, and 5 dpi, respectively.

### Inactivation of miR-183C in sensory neurons resulted in more severe reduction of CSN in the epithelial/subepithelial layer in the early stages but enhanced preservation of the large CSN in the stromal layer at later stages

Previously, we have shown that inactivation of miR-183C in CSN but not in myeloid cells results in decreased CSN density in naïve CKO vs WT control mice^40^. Consistent with this observation, at the early stage of PA infection (3 hpi), CSN density was significantly decreased in the SNS-CKO vs WT controls, in both the central and peripheral regions (**Fig.1-3**). However, CSN in CKO mice appeared to degenerate at a slower pace in the central region of the cornea in the early stage of the disease, when compared to the WT mice (**Fig.3A**). By 6 hpi, the sensory nerve density in the central region showed no significant difference between the SNS-CKO and WT control mice, because of faster degeneration in the WT mice (**Fig.3A**). Thereafter, CSN degeneration in the central region followed a similar course in both SNS-CKO and WT mice at 12 hpi, 1, 3 and 5 dpi (**Fig.2**; **Fig.3A**).

In the peripheral region of SNS-CKO mice (**Figs.2B;** orange line in **Fig.3B**), CSN degeneration followed a slower pace than the central region (orange line in **Fig.3A**). The CSN density remained relative stable in the early stages of PA infection (3 and 6 hpi)(**Fig.3B**). Similar as in the naïve mice that we reported earlier^40^, the CSN density was significantly decreased in the SNS-CKO vs WT control mice (**Figs.1, 2, 3B.** *: p<0.05). CSN density in the peripheral region did not show a significant decrease until 12 hpi in WT mice (blue line in **Fig.3B**); however, it remained stable at this time-point in the SNS-CKO mice (orange line in **Fig.3B**), erasing the difference of CSN density between SNS-CKO and WT control mice (**Fig.3B**). From 12 hpi to 1 dpi, SNS-CKO and WT mice showed similar pace of CSN degeneration in the peripheral regions of the cornea (**Fig.3B**). However, from 1 to 3 dpi, CSN density did not further decrease in the SNS-CKO mice, while in the WT mice it continued to degenerate and were nearly completely destroyed by 3 dpi (**Fig.2A-e,B-e; Fig.3B**). This resulted in higher sensory nerve density in the SNS-CKO vs WT controls at 3 dpi (**Fig.2A-e,B-e; Fig.3B**. *: p<0.05). By 5 dpi, CSN were completely degenerated in the both SNS-CKO and WT controls (**Fig.2A-f,B-f; Fig.3B**).

### Inactivation of miR-183C in the CSN resulted in changes of the dynamics of myeloid cells in the cornea during PA keratitis

To study the dynamics of myeloid cells during PA keratitis, we quantified the number of Csf1r-EGFP+ myeloid cells in the cornea at 3, 6, 12 hpi and 1, 3, and 5 dpi. At early stages of PA keratitis (3 and 6 hpi), in the WT mice, although the total number of Csf1r-EGFP+ myeloid cells per cornea showed little difference between the infected vs non-infected contralateral control eyes (**Fig.4A,B**), the density of myeloid cells in the central region of the cornea was significantly decreased, however, slightly increased in the peripheral region (**Fig.4A,C,D**. *: p<0.05). The decreased myeloid cell density in the central region is reminiscent of the macrophage/leukocyte disappearance reaction (M/LDR) that we reported early^40^.

**Figure 4.**
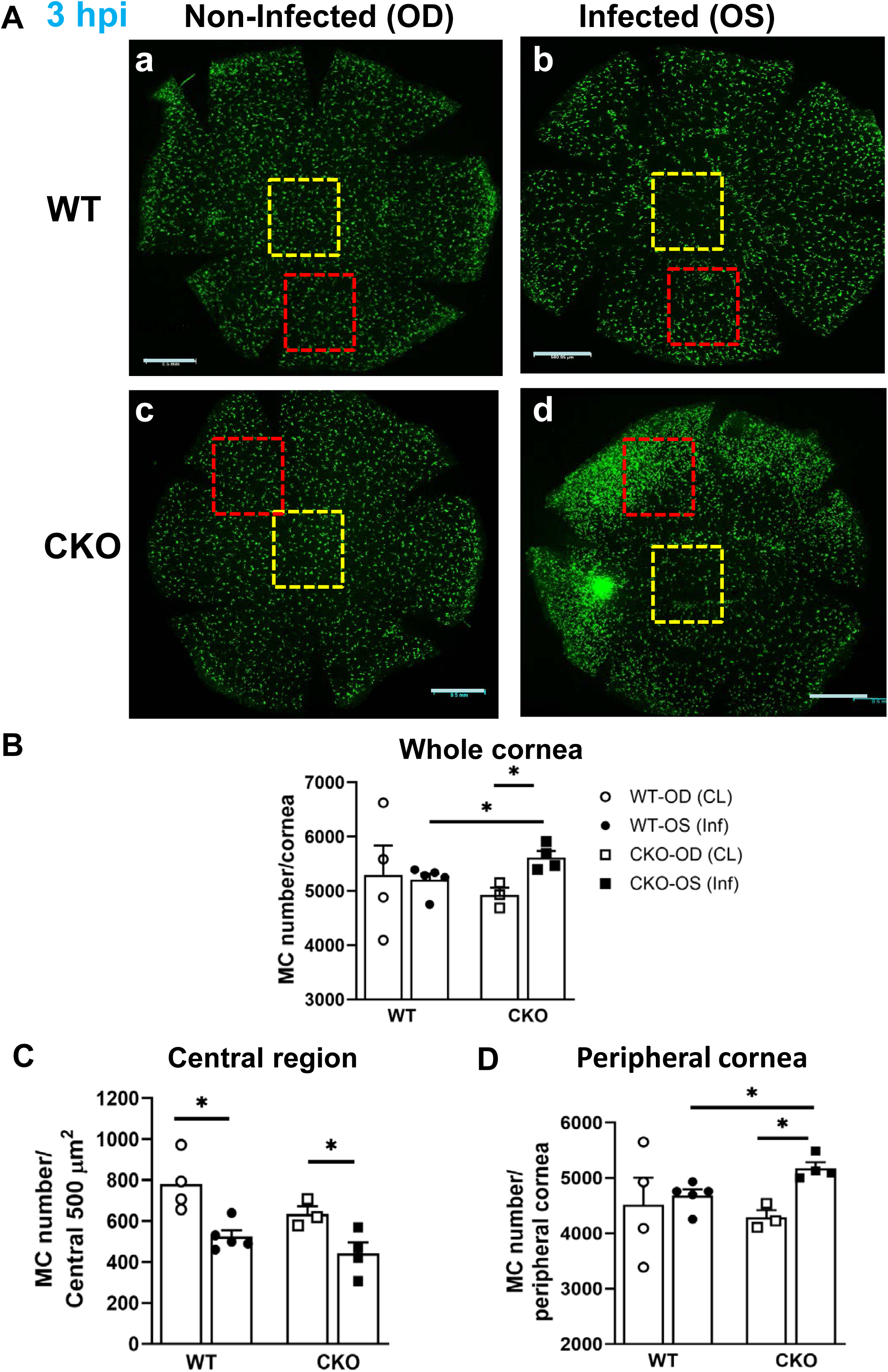
Comparison of myeloid cell number and distribution in the corneas of the SNS-CKO vs WT control mice at 3 hpi. **A**. Representatives of compressed flatmount confocal images of the infected left eyes (OS) (b,d) and non-infected contralateral eyes (OD) (a,c) of the SNS-CKO (c,d) and WT control mice (a,b). **B**. Quantification of the total number of Csf1r-EGFP+ myeloid cells (MC number)/cornea in the infected left eyes (OS, inf) and non-infected contralateral right eyes (OD, CL) of the SNS-CKO and WT control mice. **C,D**. MC number per 500 x 500 μm^2^ square in the central (C) and peripheral regions of the cornea (D).

Unlike the WT mice, in the SNS-CKO mice, the total number of myeloid cells in the cornea was increased in the infected vs non-infected contralateral eyes as early as 3 hpi, suggesting accelerated activation of immune defense machinery and faster infiltration of myeloid cells from the circulation. Densely packed Csf1r-EGFP+ myeloid cells formed infection foci (**Fig.4A-d**), which were not seen in the WT control mice at this stage (**Fig.4A-b**), suggesting accelerated infiltration in the SNS-CKO mice. In spite of the enhanced infiltration of myeloid cells in the peripheral regions of the cornea, like in the WT mice, the density of myeloid cells in the central region of the infected vs non-infected, contralateral control cornea was also decreased in the SNS-CKO mice (**Fig.4A,B**), suggesting the M/LDR was not affected by SNS-CKO of miR-183C, consistent with our previous report^40^.

At 6 hpi, myeloid cells in the infected eyes of SNS-CKO mice remained at a similar level as 3 hpi, while the number of myeloid cells in the corneas of infected eyes of WT mice was further increased to a similar level as the SNS-CKO mice (**Fig.5A,B**). Similar to the SNS-CKO mice, densely packed infection foci appeared in the infected corneas of the WT mice (**Fig.5A-b**), although the sizes of foci appeared to be smaller than in the SNS-CKO mice (**Fig.5A-b&d**).

**Figure 5.**
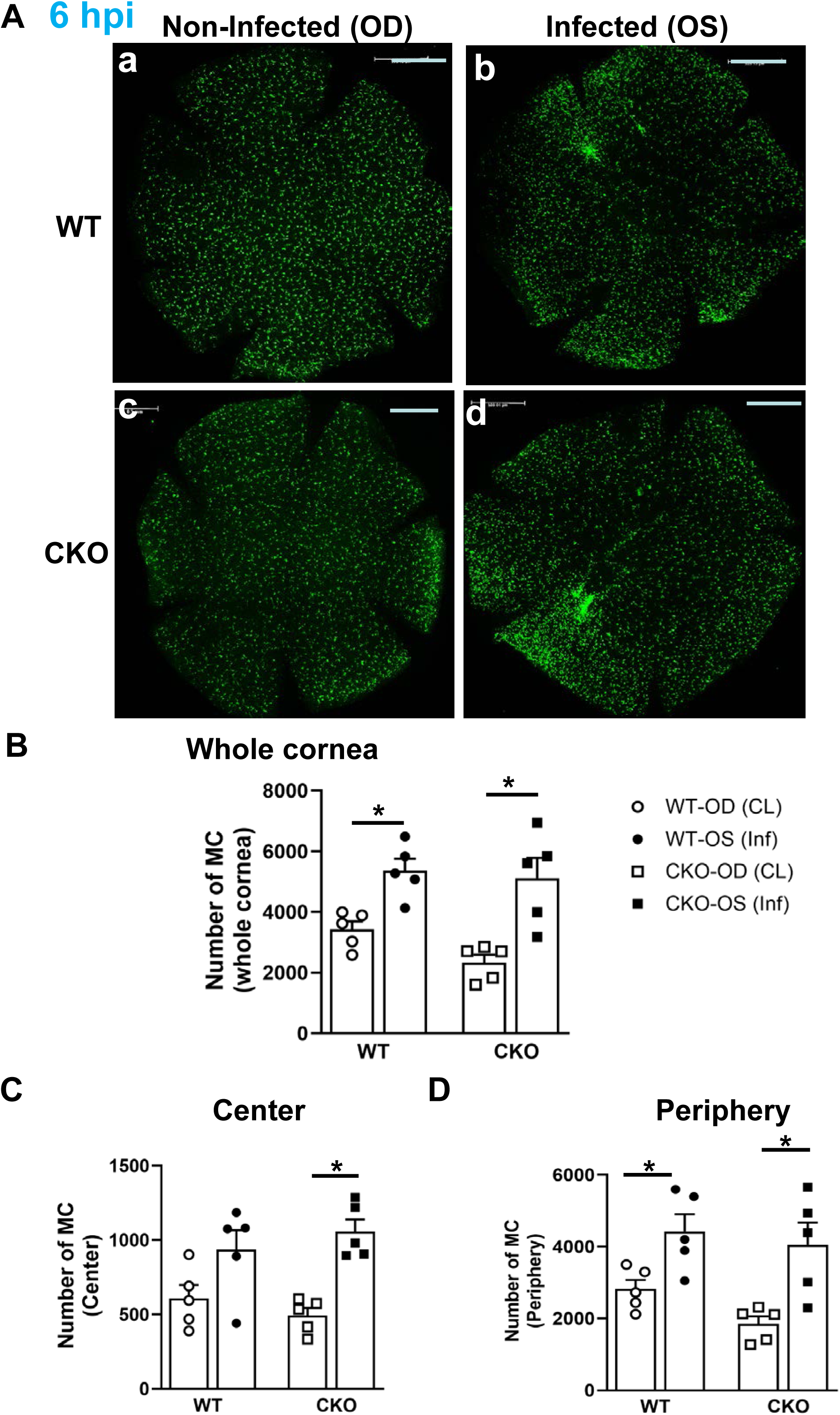
Comparison of myeloid cell number and distribution in the corneas of the SNS-CKO vs WT control mice at 6 hpi. **A**. Representatives of compressed flatmount confocal images of the infected left eyes (OS) (b,d) and non-infected contralateral eyes (OD) (a,c) of the SNS-CKO (c,d) and WT control mice (a,b). Scale bars: 500 μm. **B**. Quantification of the total number of Csf1r-EGFP+ myeloid cells (MC number)/cornea. **C,D**. MC number per 500 x 500 μm^2^ square in the central (C) and peripheral regions of the cornea (D).

From 6 to 12 hpi, drastic change occurred in the infected corneas (**Fig.6**). At 12 hpi, in both SNS-CKO and WT mice, densely-packed myeloid cells formed a ring-shaped band circling a “dark zone” in the center of the cornea, which was nearly devoid of myeloid cells, and appeared to coincide with the central area of the cornea where CSN had degenerated (yellow circles in **Fig.6A**). Since it is impossible to count the number of EGFP+ myeloid cells, we quantified the volume of the EGFP fluorescence to reflect the density of myeloid cells between SNS-CKO and WT controls. Our data showed that at 12 hpi, although the overall of intensity of EGFP signals showed no difference between SNS-CKO, vs WT controls (**Fig.6B**), the size of the central “dark zone” was larger in the SNS-CKO vs WT mice, consistent with enhanced CSN degeneration (**Fig.6A,C**), while the density of myeloid cells in the central “dark zone” were decreased in SNS-CKO vs WT controls (**Fig.6A,D**. *: p<0.05).

**Figure 6.**
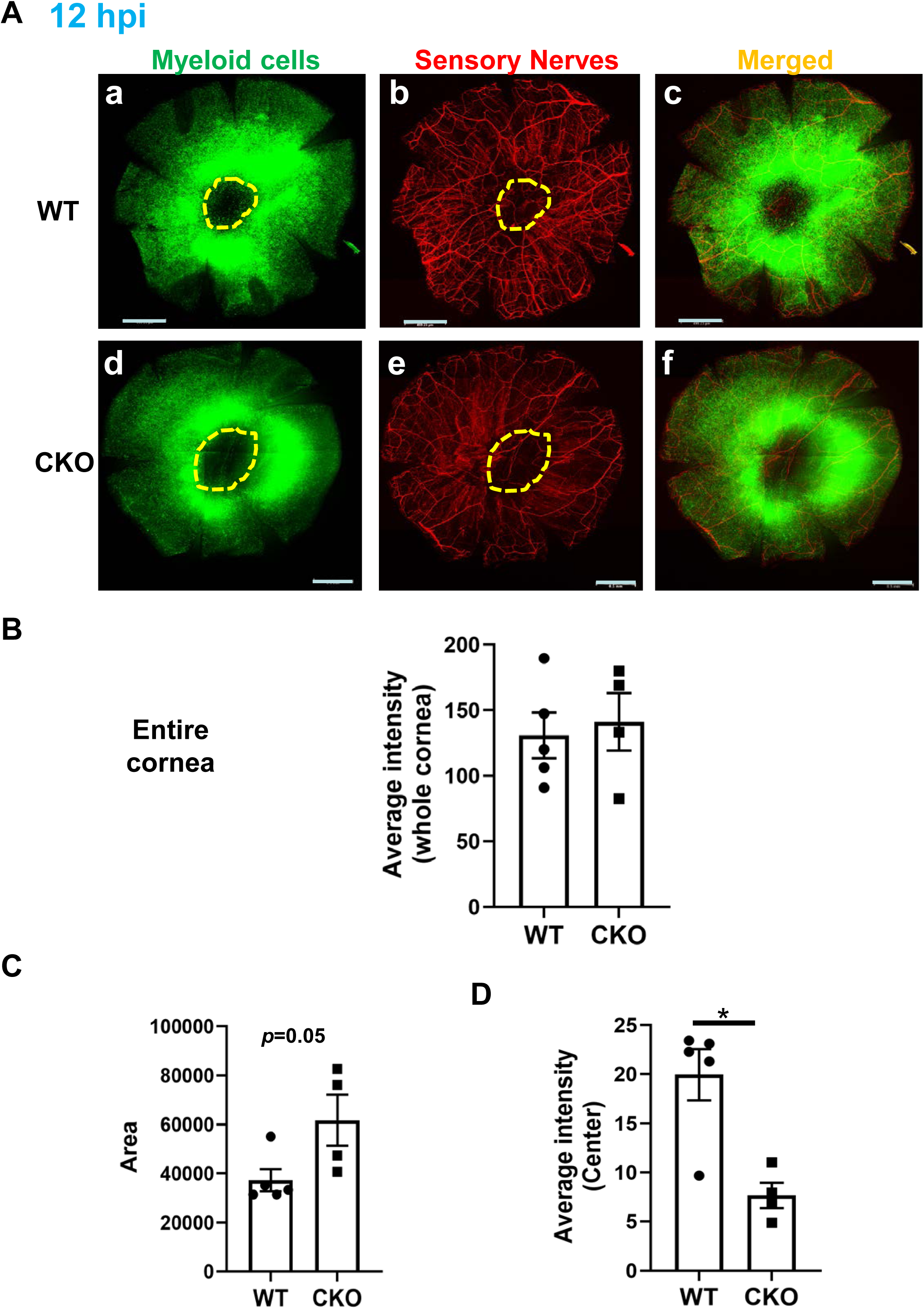
Comparison of myeloid cell distribution in relation to CSN in the corneas of the SNS-CKO vs WT control mice at 12 hpi. **A**. Representatives of compressed flatmount confocal images of the myeloid cells (GFP+, a,d), sensory nerves (RFP+, b,e) and merged images (c,f) of the corneas of infected eyes of the SNS-CKO (d-f) and WT control mice (a-c). Scale bars: 500 μm. **B**. Quantification of the average intensity of EGFP fluorescence of the whole cornea. **C,D.** The area (C) and average intensity of the EGFP fluorescence of the MC (D) in the “dark zone” encircled by the inflammatory ring. Scale bars: 500 μm.

At 1 dpi, the entire corneas appeared to be filled with densely packed myeloid cells in both SNS-CKO and WT control mice (**Fig.7**). However, the average intensity of EGFP in the CKO mice was decreased (**Fig.7B**) and the area filled with EGFP+ myeloid cells was reduced (**Fig.7C**), when compared to the WT mice, suggesting decreased infiltration of myeloid cells in the cornea.

**Figure 7.**
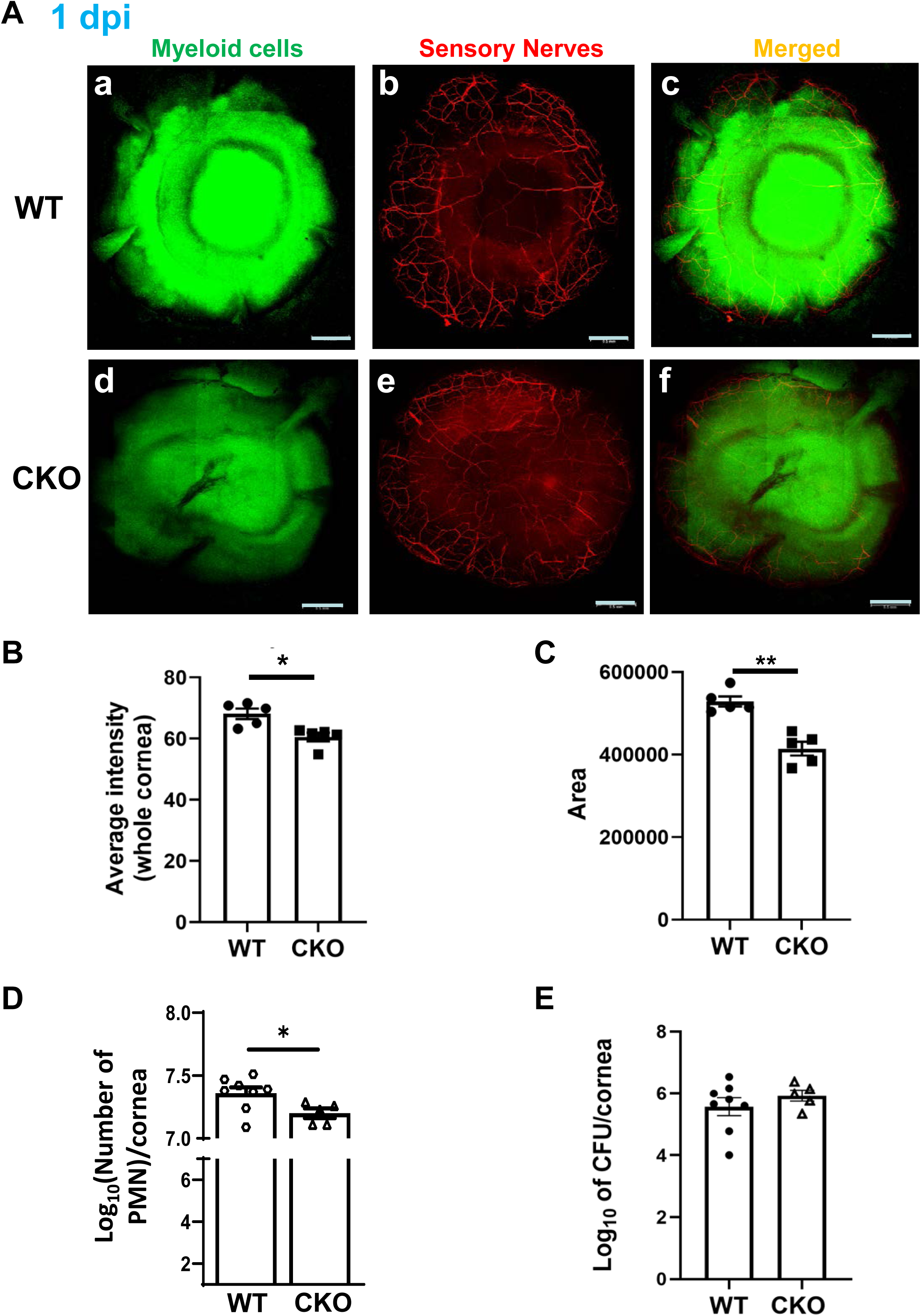
Comparison of myeloid cell distribution in relation to CSN in the corneas of the SNS-CKO vs WT control mice at 1 dpi. **A**. Representatives of compressed flatmount confocal images of the myeloid cells (GFP+, a,d), CSN (RFP+, b,e) and merged images (c,f) of the corneas of infected eyes of the SNS-CKO (d-f) and WT control mice (a-c). Scale bars: 500 μm. **B,C**. Quantification of the average intensity (B) and the area of EGFP fluorescence of the whole cornea (C). **D,E.** The number of PMN (D) and residual bacteria per cornea at 1 dpi (E).

To test this, we performed MPO assay to quantify the number of neutrophils in the infected cornea. Our result showed that the number of neutrophils in the SNS-CKO mice was indeed significantly decreased (∼1.4 fold) when compared to the WT mice at 1 dpi (**Fig.7D**). It is estimated that the SNS-CKO mice has ∼7 million less neutrophils/cornea than the WT, based that 1 unit of MPO is equivalent to approximately 2x10^5^ neutrophils^53^. Consistently, a viable bacterial plate assay to quantify the residual bacteria in the infected cornea of SNS-CKO mice showed slightly increased number of colony forming units (CFU) of PA, when compare to the WT controls (∼2.3 fold increase and ∼ 4.7 x 10^5^ more bacteria/cornea), although the data did not pass the threshold of statistical significance (**Fig.7E**).

From 3 dpi (**Fig.8A-B**), the entire cornea was filled with EGFP+ myeloid cells. No quantitative difference between WT and CKO was observed.

**Figure 8.**
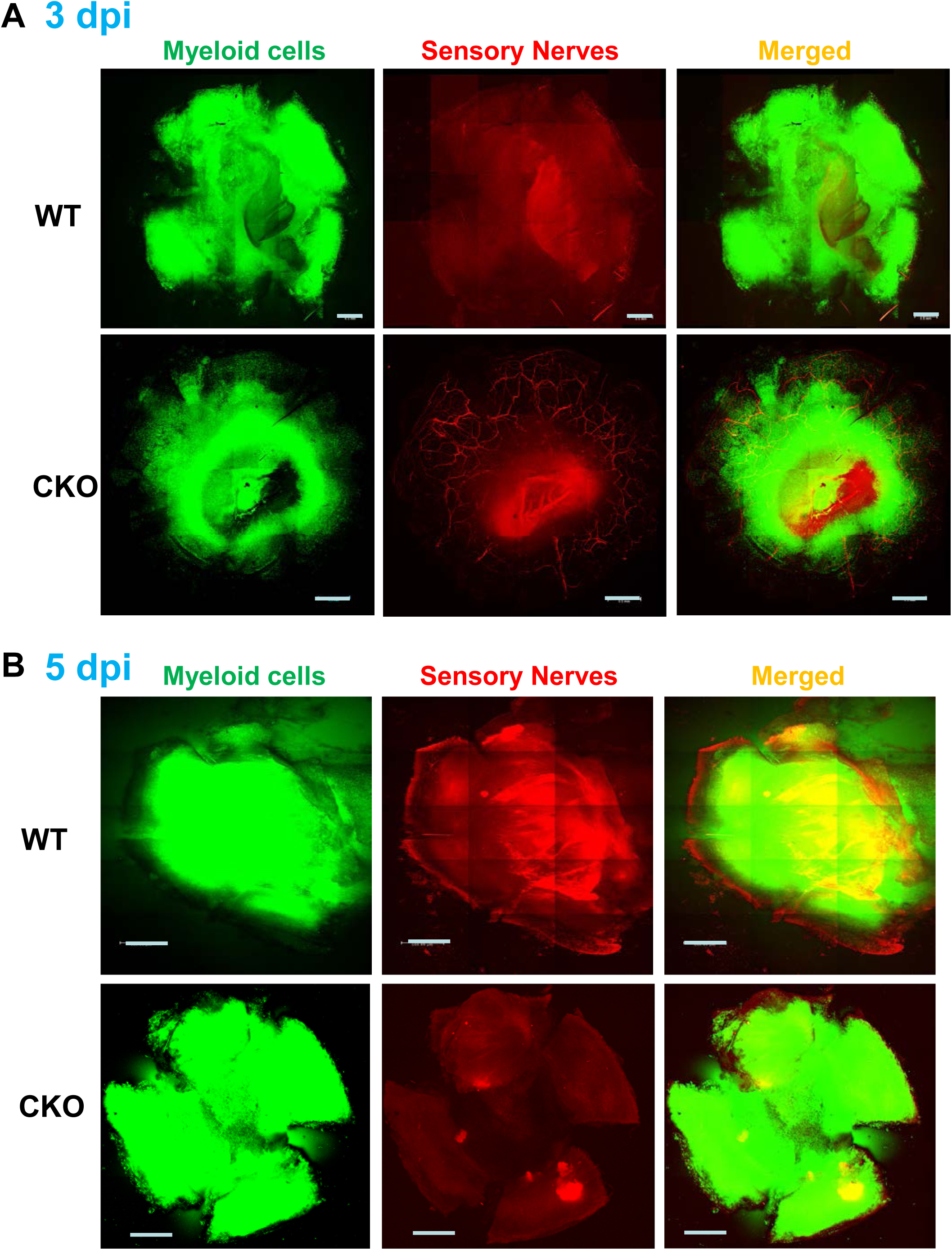
Myeloid cell distribution in relation to CSN in the corneas of the SNS-CKO vs WT control mice at 3 dpi (A) and 5 dpi (B).

### Inactivation of miR-183C in the sensory neurons resulted in a decrease of the severity of PA keratitis at 3 dpi

In spite the decreased neutrophil infiltration in the SNS-CKO vs WT mice at 1 dpi, the disease severity did not show significant difference at this time-point (**Fig.9**). However, at 3 dpi, PA keratitis showed a slight decreased severity in the SNS-CKO mice (**Fig.9**). At this stage, the infected corneas of 8 out of 9 WT mice were perforated; while only 4 out of 10 SNS-CKO mice were perforated; the clinical scores of the rest 6 SNS-CKO remained at +3 (**Fig.9**). At 5 dpi, no significant difference in clinical scores was observed between the SNS-CKO and WT control mice (**Fig.9**).

**Figure 9.**
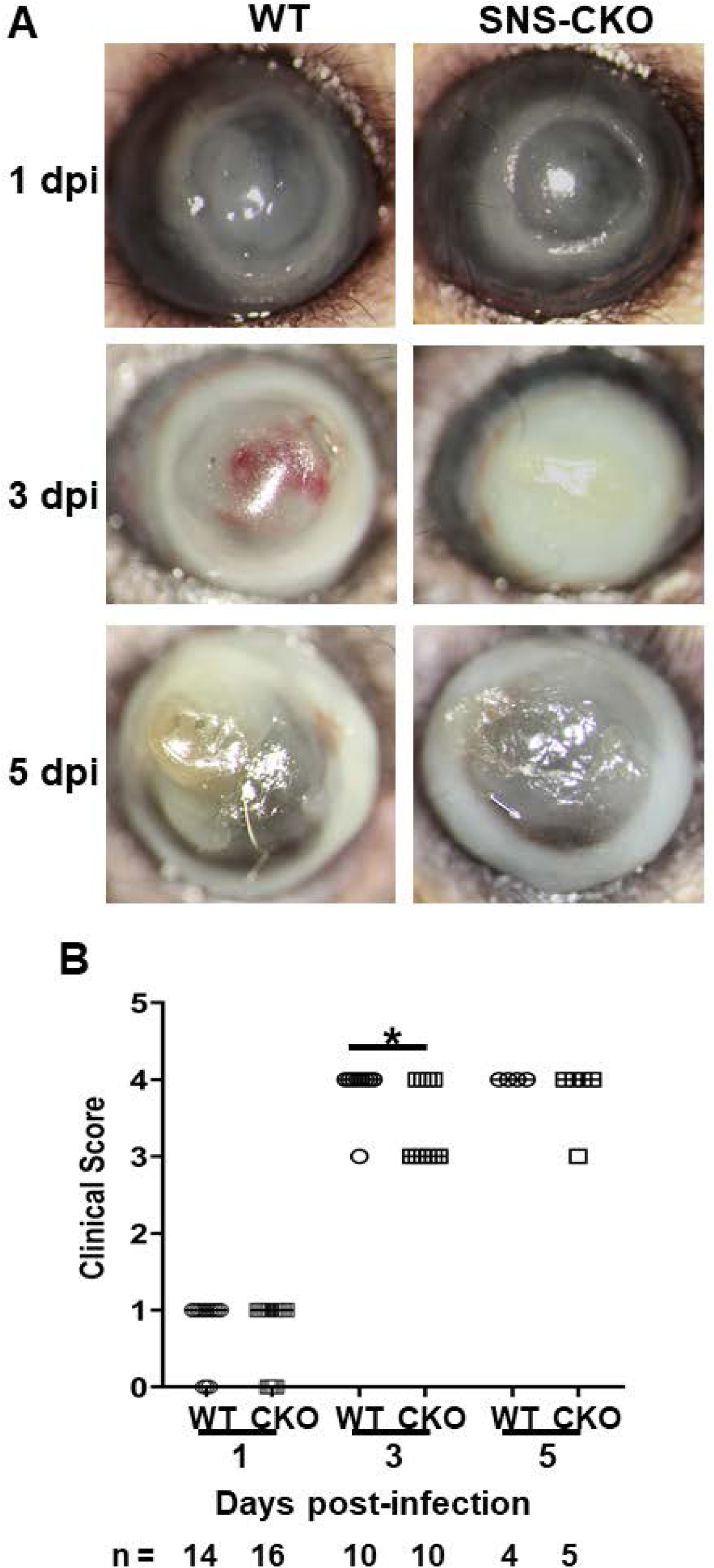
SNS-CKO of miR-183C resulted in slightly decreased severity of PA keratitis at 3 dpi. **A**. Representative slit lamp photographs of SNS-CKO and WT control mice at 1, 3 and 5 dpi. **B**. Clinical scores. n numbers at each time-point and genotype are specified at the bottom.

## Discussion and Conclusion

The miR-183C is required for the normal functions of both CSN and innate immune cells^39–42^. Complete inactivation of miR-183C in mice has significant functional impacts on both corneal sensory innervation and innate immunity, resulting in decreased CSN density and a disrupted structural pattern of the subbasal plexus of the cornea^39^. It also reduces pro-inflammatory responses while enhancing the phagocytic and bacterial killing capacity of innate immune cells, including Mφ and neutrophils^39, 42^. These lead to an overall decreased severity of PA keratitis in the miR-183C conventional KO vs WT control mice^39^. Consistently, corneal knockdown of miR-183C by anti-miR-183C application to the cornea showed similar effect^43^, suggesting miR-183C modulates PA keratitis through its dual regulation of CSN and innate immunity. Here, to further specify the contribution of its regulation of sensory innervation to the modulation of PA keratitis, we created a SNS-CKO model. In comparison to their WT controls, our data showed that inactivation of SNS-CKO of miR-183C resulted in changes in the dynamics of sensory nerve degeneration (intrinsic regulation) as well as innate immune cell infiltration in the cornea (extrinsic regulation) - a display of the effect of neuroimmune interactions during PA keratitis. The severity of the disease in the SNS-CKO vs WT control mice was only modestly decreased when compared to what we observed in the conventional KO mice^39^ or acute miR-183C knockdown^43^, in which miR-183C is inactivated^39^ or down-regulated in both sensory nerves and innate immune cells^43^. This observation suggests that effects of knockout/knockdown of miR-183C in sensory nerves and innate immunity are additive, if not synergistic; in combination, they result in more robust decrease of the severity of PA keratitis. Simultaneous knockdown of miR-183C in CSN and myeloid cells is required for a therapeutic approach. A myeloid cell-specific CKO of miR-183C mouse model will help to further elucidate whether and how miR-183C’s functions in sensory nerves and innate immune cells work additively or synergistically in modulation of PA keratitis.

Although it has been reported that CSN are destroyed quickly after PA infection^14, 20^, previous studies were conducted by immunostaining of pan-neuronal marker, β-III tubulin, and did not provide clear images of the entire cornea with spatial and temporal details^14, 20^. Therefore, the behavior of the sensory innervation during PA keratitis remain unresolved. In the current study, the double tracing mouse model, in which corneal sensory nerves are labeled with Nav1.8-Cre driven RFP reporter, while myeloid cells are labeled with Csf1r-EGFP, provides a powerful tool to simultaneously observe the behaviors and sensory nerves and innate immune cells and their interactions during PA keratitis with unprecedented resolution. Our study provides new insights into the dynamic changes of CSN during PA keratitis with spatial and temporal details. We showed that CSN degeneration began as early as 3 hpi, starting with fine terminal nerves in the epithelial/subepithelial layers in the central region of the cornea. The CSN degeneration gradually progressed to the periphery of the cornea. By 1 dpi, sensory nerves in the epithelial layer were nearly completely destroyed, however, large-diameter nerves and their branches in the stromal layer persisted. Since CSN modulate corneal immune/inflammatory responses through exocytosis at their nerve endings, which are mostly distributed in the epithelial and subepithelial plexus^12, 54, 55^, based on our result, it is reasonable to speculate that modulation of the immune response to PA infection by the sensory nerves occurs mostly in the first 24-hour window. However, we cannot exclude the possibility that sensory nerves remodel their synaptic distribution during PA keratitis and modulate the disease process in later stages.

Previously, we and other have shown that miR-183C is responsible for the terminal functional differentiation and organization in sensory neurons of various sensory organs^38, 56–59^.

Consistently, complete inactivation or knockdown as well as SNS-CKO of miR-183C result in decreased sensory serve density and interrupted patterns of the fine terminal nerves in the epithelial and subbasal plexus of the cornea of naïve KO or SNS-CKO mice when compared to their corresponding WT controls^39, 40, 43^. This suggests that miR-183C promotes corneal sensory neurite growth and patterning during corneal development and the establishment of the CSN innervation. This effect is a result of its direct regulation of the molecules involved in axon guidance and neuronal projection-related genes^40, 43^. Our observation of decreased CSN density in the SNS-CKO mice in the early stage of the PA keratitis is possibly a reflection of this effect.

However, our data also suggest that the CSN showed an accelerated degeneration in the WT mice in the early stage of PA keratitis; by 6 hpi, the difference in the sensory nerve density between the SNS-CKO and WT controls were “erased”. At 3 dpi, the residual large sensory nerves were better preserved in the SNS-CKO vs WT control mice. These results support a hypothesis that, in the context of PA keratitis, downregulation of miR-183C may have a protective effect on the large-diameter nerves in the stroma. Consistent with this hypothesis, our previously published transcriptomic data suggests that, in addition to neuron projection development-related genes, miR-183C regulates neuronal survival/apoptosis through targeting a series of key molecules in these processes^40^. Future studies will be needed to directly test this hypothesis.

Besides the behavior of CSN, our study also provides new insights into the dynamics of innate immune cells during PA keratitis. Here we showed that at the early stages of the disease (3 and 6 hpi), the density of myeloid cells was decreased in the central region of the cornea, while increased in the peripheral region. The decrease in the myeloid cell density in the central region is possibly a result of the M/LDR that we reported earlier^41^; while the increased density in the peripheral region of the cornea a result of infiltration. M/LDR is considered to allow the immune-regulatory resident Mφ to temporarily “give way” to pro-inflammatory myeloid cells to evoke an acute inflammatory response before they “re-appear” to contain the inflammation and promote tissue repair ^60–62^. Our data supports this hypothesis. However, the current mouse model cannot distinguish between the resident myeloid cells from the infiltrating cells. Future studies with specific lineage tracing capability will further elucidate this.

Furthermore, our data, for the first time, simultaneously revealed the dynamic changes of CSN and myeloid cells during PA keratitis, providing an opportunity to uncover the interactions of CSN and innate immune cells. Consistent with our previous report^41^, our data showed that in the early stage of PA keratitis (3 and 6 hpi), infiltration of myeloid cells was accelerated in the SNS-CKO vs WT control mice, suggesting an extrinsic regulation of miR-183C in CSN of the innate immune responses. It is possibly a result of the direct regulations of miR-183C on pro-inflammatory neuropeptides, including Cx3cl1, which is known to promote chemotactic migration of microglial in the central nervous system^63, 64^, and mediate the homing of resident myeloid cells in the cornea^65^ and recruitment of Mφ in other tissues^66^. Additional studies will be needed to further test this hypothesis.

At 12 dpi, the densely-packed infiltrating myeloid cells formed a ring-shaped band, which we designate here as an “inflammatory ring”. This inflammatory ring circled a zone in the central region of the cornea, which was nearly devoid of myeloid cells, coinciding with the central region of the cornea where CSN degenerated first. The number of myeloid cells was decreased in the ring center of the SNS-CKO vs WT controls, while CSN density was reduced, suggesting a contribution of CSN degeneration in the formation of the “inflammatory ring” – neuroimmune interaction. The inflammatory ring appeared to be a transient phenomenon, as, by 1 dpi, the infected cornea including the central region was filled with myeloid cells. The functional significance of the inflammatory ring during PA keratitis warrants further studies. It is well known that different mouse strains with different genetic background have their differences in corneal nerve densities^67^ and immune responses to PA infection^68–71^. In the current study, all mice were on the C57BL/6 background, which is known to favor Th1 responsiveness and has enhanced susceptibility and severe PA keratitis^69^. Mice with different genetic backgrounds are known to have different immune/inflammatory responses in PA keratitis. For example, mouse strains with Th2-dominant response, e.g., BALB/c, are relatively resistant to PA infection and have a milder course of disease without corneal perforation^68–71^. Additional studies are required to determine whether our observations of the neuroimmune responses here apply to other mouse models and as well as in humans.

A few limitations of the current study need to be addressed in our future endeavors. First, although the flatmount microscopic study illustrated greater details of the simultaneous changes of CSN and myeloid cells during PA keratitis, they are static images of neuro-immune components at different stages of the disease. Time-lapse live imaging by advanced multiphoton microscopy will be required to provide further insights of neuroimmune interactions. Second, in this study, we focused on the interaction of sensory neurons and innate immune cells. However, increasing evidence suggests corneal sympathetic nerves interact with CSN,

CRICs and other cellular components and modulate corneal homeostasis and pathogenesis of various corneal diseases. For example, the sympathetic nerves have been shown to promote inflammatory responses and delay corneal re-epithelialization after epithelial debridement^72–74^. Inhibition of sympathetic innervation alleviated acute tobacco smoke induced deficiencies in corneal wound healing^75^. In herpes stromal keratitis (HSK), it has been shown that CD4+ T cell-and myeloid cell-produced VEGF causes sensory nerve regression and simultaneously enhances sympathetic innervation in the cornea^76^. The sympathetic neurotransmitter, norepinephrine (NE), has been shown to aggravate bacterial keratitis by a combination of compromising epithelial integrity, enhancing bacterial growth and virulence, and inflammatory responses^77–79^. In addition, our recent systems biology approach using single-cell RNA sequencing technology showed that miR-183C’s regulation is not restricted to the immune cells and sensory nerves, but also imposes global modulation of the entire corneal cellular landscape^80^. Studies of the simultaneous changes and interactions among CSN, sympathetic nerves, innate immunes and other components of the cornea with a systems biology approach are warranted.

Despite limitations, our current study created a powerful model to study (sensory) neuroimmune interactions during PA keratitis and provided new insights into the molecular mechanisms of PA keratitis, paving the way to future endeavors to fully elucidate the molecular mechanisms of the pathogenesis of PA keratitis.

## Acknowledgement

This work is supported by grants from the National Eye Institute, National Institutes of Health (R01 EY026059 to SX; R01 EY035231, and P30 EY004068 to LDH); a Research to Prevent Blindness unrestricted grant to the Department of Ophthalmology, Visual and Anatomical Science, Wayne State University School of Medicine.

## References

1. Hazlett LD. Corneal response to Pseudomonas aeruginosa infection. Prog Retin Eye Res 2004;23:1–30.

2. Choy MH, Stapleton F, Willcox MD, Zhu H. Comparison of virulence factors in Pseudomonas aeruginosa strains isolated from contact lens-and non-contact lens-related keratitis. J Med Microbiol 2008;57:1539–1546.

3. Stapleton F, Carnt N. Contact lens-related microbial keratitis: how have epidemiology and genetics helped us with pathogenesis and prophylaxis. Eye (Lond) 2012;26:185–193.

4. Green M, Apel A, Stapleton F. Risk factors and causative organisms in microbial keratitis. Cornea 2008;27:22–27.

5. Tam C, Mun JJ, Evans DJ, Fleiszig SM. The impact of inoculation parameters on the pathogenesis of contact lens-related infectious keratitis. Invest Ophthalmol Vis Sci 2010;51:3100–3106.

6. Gadjeva M, Nagashima J, Zaidi T, Mitchell RA, Pier GB. Inhibition of macrophage migration inhibitory factor ameliorates ocular Pseudomonas aeruginosa-induced keratitis. PLoS Pathog 2010;6:e1000826.

7. O’Brien TP, Maguire MG, Fink NE, Alfonso E, McDonnell P. Efficacy of ofloxacin vs cefazolin and tobramycin in the therapy for bacterial keratitis. Report from the Bacterial Keratitis Study Research Group. Arch Ophthalmol 1995;113:1257–1265.

8. Mesaros N, Nordmann P, Plesiat P, et al. Pseudomonas aeruginosa: resistance and therapeutic options at the turn of the new millennium. Clin Microbiol Infect 2007;13:560–578.

9. Ung L, Acharya NR, Agarwal T, et al. Infectious corneal ulceration: a proposal for neglected tropical disease status. Bull World Health Organ 2019;97:854–856.

10. Chodosh J. Infectious corneal ulceration (ICU)-an enduring unmet need., ARVO2021. Virtual meeting; 2021.

11. Marfurt CF, Cox J, Deek S, Dvorscak L. Anatomy of the human corneal innervation. Exp Eye Res 2010;90:478–492.

12. Muller LJ, Marfurt CF, Kruse F, Tervo TM. Corneal nerves: structure, contents and function. Exp Eye Res 2003;76:521–542.

13. Guerrero-Moreno A, Baudouin C, Melik Parsadaniantz S, Reaux-Le Goazigo A. Morphological and Functional Changes of Corneal Nerves and Their Contribution to Peripheral and Central Sensory Abnormalities. Front Cell Neurosci 2020;14:610342.

14. Lin T, Quellier D, Lamb J, et al. Pseudomonas aeruginosa-induced nociceptor activation increases susceptibility to infection. PLoS Pathog 2021;17:e1009557.

15. Zhou Z, Barrett RP, McClellan SA, et al. Substance P delays apoptosis, enhancing keratitis after Pseudomonas aeruginosa infection. Invest Ophthalmol Vis Sci 2008;49:4458–4467.

16. Lighvani S, Huang X, Trivedi PP, Swanborg RH, Hazlett LD. Substance P regulates natural killer cell interferon-gamma production and resistance to Pseudomonas aeruginosa infection. Eur J Immunol 2005;35:1567–1575.

17. Hazlett LD, McClellan SA, Barrett RP, Liu J, Zhang Y, Lighvani S. Spantide I decreases type I cytokines, enhances IL-10, and reduces corneal perforation in susceptible mice after Pseudomonas aeruginosa infection. Invest Ophthalmol Vis Sci 2007;48:797–807.

18. McClellan SA, Zhang Y, Barrett RP, Hazlett LD. Substance P promotes susceptibility to Pseudomonas aeruginosa keratitis in resistant mice: anti-inflammatory mediators downregulated. Invest Ophthalmol Vis Sci 2008;49:1502–1511.

19. Foldenauer ME, McClellan SA, Barrett RP, Zhang Y, Hazlett LD. Substance P affects growth factors in Pseudomonas aeruginosa-infected mouse cornea. Cornea 2012;31:1176–1188.

20. Yuan K, Zheng J, Shen X, et al. Sensory nerves promote corneal inflammation resolution via CGRP mediated transformation of macrophages to the M2 phenotype through the PI3K/AKT signaling pathway. Int Immunopharmacol 2022;102:108426.

21. Ambros V. The functions of animal microRNAs. Nature 2004;431:350–355.

22. Bartel DP. MicroRNAs: genomics, biogenesis, mechanism, and function. Cell 2004;116:281–297.

23. Wightman B, Ha I, Ruvkun G. Posttranscriptional regulation of the heterochronic gene lin-14 by lin-4 mediates temporal pattern formation in C. elegans. Cell 1993;75:855–862.

24. Lee RC, Feinbaum RL, Ambros V. The C. elegans heterochronic gene lin-4 encodes small RNAs with antisense complementarity to lin-14. Cell 1993;75:843–854.

25. Chang TC, Mendell JT. microRNAs in vertebrate physiology and human disease. Annu Rev Genomics Hum Genet 2007;8:215–239.

26. Alvarez-Garcia I, Miska EA. MicroRNA functions in animal development and human disease. Development 2005;132:4653–4662.

27. Lee YS, Dutta A. MicroRNAs in cancer. Annu Rev Pathol 2009;4:199–227.

28. Mencia A, Modamio-Hoybjor S, Morin M, et al. A mutation in the human miR-96, a microRNA expressed in the inner ear, causes non-syndromic progressive hearing loss. 6th Molecular Biology of Hearing and Hearing Deafness Conference. Wellcome Trust Conference Center, Hinxton, UK; 2007.

29. Hughes AE, Bradley DT, Campbell M, et al. Mutation altering the miR-184 seed region causes familial keratoconus with cataract. Am J Hum Genet 2011;89:628–633.

30. Iliff BW, Riazuddin SA, Gottsch JD. A single-base substitution in the seed region of miR-184 causes EDICT syndrome. Invest Ophthalmol Vis Sci 2012;53:348–353.

31. Conte I, Hadfield KD, Barbato S, et al. MiR-204 is responsible for inherited retinal dystrophy associated with ocular coloboma. Proc Natl Acad Sci U S A 2015.

32. Xu S. microRNAs and inherited retinal dystrophies. Proc Natl Acad Sci U S A 2015;112:8805–8806.

33. Jeyaseelan K, Herath WB, Armugam A. MicroRNAs as therapeutic targets in human diseases. Expert Opin Ther Targets 2007;11:1119–1129.

34. van Rooij E, Olson EN. MicroRNAs: powerful new regulators of heart disease and provocative therapeutic targets. J Clin Invest 2007;117:2369–2376.

35. Zhang B, Farwell MA. microRNAs: a new emerging class of players for disease diagnostics and gene therapy. J Cell Mol Med 2008;12:3–21.

36. Stenvang J, Kauppinen S. MicroRNAs as targets for antisense-based therapeutics. Expert Opin Biol Ther 2008;8:59–81.

37. Xu S, Witmer PD, Lumayag S, Kovacs B, Valle D. MicroRNA (miRNA) Transcriptome of Mouse Retina and Identification of a Sensory Organ-specific miRNA Cluster. J Biol Chem 2007;282:25053–25066.

38. Lumayag S, Haldin CE, Corbett NJ, et al. Inactivation of the microRNA-183/96/182 cluster results in syndromic retinal degeneration. Proc Natl Acad Sci U S A 2013;110:E507–516.

39. Muraleedharan CK, McClellan SA, Barrett RP, et al. Inactivation of the miR-183/96/182 Cluster Decreases the Severity of Pseudomonas aeruginosa-Induced Keratitis. Invest Ophthalmol Vis Sci 2016;57:1506–1517.

40. Gupta N, Somayajulu M, Gurdziel K, et al. The miR-183/96/182 cluster regulates sensory innervation, resident myeloid cells and functions of the cornea through cell type-specific target genes. Scientific reports 2024;14:7676.

41. Coku A, McClellan SA, Van Buren E, Back JB, Hazlett LD, Xu S. The miR-183/96/182 Cluster Regulates the Functions of Corneal Resident Macrophages. Immunohorizons 2020;4:729–744.

42. Muraleedharan CK, McClellan SA, Ekanayaka SA, et al. The miR-183/96/182 Cluster Regulates Macrophage Functions in Response to Pseudomonas aeruginosa. J Innate Immun 2019;1–12.

43. McClellan S, Pitchaikannu A, Wright R, et al. Prophylactic Knockdown of the miR-183/96/182 Cluster Ameliorates Pseudomonas aeruginosa-Induced Keratitis. Invest Ophthalmol Vis Sci 2021;62:14.

44. Peng C, Li L, Zhang MD, et al. miR-183 cluster scales mechanical pain sensitivity by regulating basal and neuropathic pain genes. Science 2017;356:1168–1171.

45. Stirling LC, Forlani G, Baker MD, et al. Nociceptor-specific gene deletion using heterozygous NaV1.8-Cre recombinase mice. Pain 2005;113:27–36.

46. Kozak CA, Sangameswaran L. Genetic mapping of the peripheral sodium channel genes, Scn9a and Scn10a, in the mouse. Mamm Genome 1996;7:787–788.

47. Souslova VA, Fox M, Wood JN, Akopian AN. Cloning and characterization of a mouse sensory neuron tetrodotoxin-resistant voltage-gated sodium channel gene, Scn10a. Genomics 1997;41:201–209.

48. Abrahamsen B, Zhao J, Asante CO, et al. The cell and molecular basis of mechanical, cold, and inflammatory pain. Science 2008;321:702–705.

49. Agarwal N, Offermanns S, Kuner R. Conditional gene deletion in primary nociceptive neurons of trigeminal ganglia and dorsal root ganglia. Genesis 2004;38:122–129.

50. Sasmono RT, Oceandy D, Pollard JW, et al. A macrophage colony-stimulating factor receptor-green fluorescent protein transgene is expressed throughout the mononuclear phagocyte system of the mouse. Blood 2003;101:1155–1163.

51. MacDonald KP, Rowe V, Bofinger HM, et al. The colony-stimulating factor 1 receptor is expressed on dendritic cells during differentiation and regulates their expansion. J Immunol 2005;175:1399–1405.

52. Madisen L, Zwingman TA, Sunkin SM, et al. A robust and high-throughput Cre reporting and characterization system for the whole mouse brain. Nat Neurosci 2010;13:133–140.

53. Williams RN, Paterson CA, Eakins KE, Bhattacherjee P. Quantification of ocular inflammation: evaluation of polymorphonuclear leucocyte infiltration by measuring myeloperoxidase activity. Curr Eye Res 1982;2:465–470.

54. Hwang DD, Lee SJ, Kim JH, Lee SM. The Role of Neuropeptides in Pathogenesis of Dry Dye. J Clin Med 2021;10.

55. Liu J, Huang S, Yu R, et al. TRPV1(+) sensory nerves modulate corneal inflammation after epithelial abrasion via RAMP1 and SSTR5 signaling. Mucosal Immunol 2022;15:867–881.

56. Geng R, Furness DN, Muraleedharan CK, et al. The microRNA-183/96/182 Cluster is Essential for Stereociliary Bundle Formation and Function of Cochlear Sensory Hair Cells. Scientific reports 2018;8:18022.

57. Kuhn S, Johnson SL, Furness DN, et al. miR-96 regulates the progression of differentiation in mammalian cochlear inner and outer hair cells. Proc Natl Acad Sci U S A 2011;108:2355–2360.

58. Fan J, Jia L, Li Y, et al. Maturation arrest in early postnatal sensory receptors by deletion of the miR-183/96/182 cluster in mouse. Proc Natl Acad Sci U S A 2017;114:E4271–E4280.

59. Xiang L, Chen XJ, Wu KC, et al. miR-183/96 plays a pivotal regulatory role in mouse photoreceptor maturation and maintenance. Proc Natl Acad Sci U S A 2017;114:6376–6381.

60. Barth MW, Hendrzak JA, Melnicoff MJ, Morahan PS. Review of the macrophage disappearance reaction. J Leukoc Biol 1995;57:361–367.

61. Davies LC, Rosas M, Jenkins SJ, et al. Distinct bone marrow-derived and tissue-resident macrophage lineages proliferate at key stages during inflammation. Nat Commun 2013;4:1886.

62. Liu J, Xue Y, Dong D, et al. CCR2(-) and CCR2(+) corneal macrophages exhibit distinct characteristics and balance inflammatory responses after epithelial abrasion. Mucosal Immunol 2017;10:1145–1159.

63. Harrison JK, Jiang Y, Chen S, et al. Role for neuronally derived fractalkine in mediating interactions between neurons and CX3CR1-expressing microglia. Proc Natl Acad Sci U S A 1998;95:10896–10901.

64. Pawelec P, Ziemka-Nalecz M, Sypecka J, Zalewska T. The Impact of the CX3CL1/CX3CR1 Axis in Neurological Disorders. Cells 2020;9.

65. Chinnery HR, Ruitenberg MJ, Plant GW, Pearlman E, Jung S, McMenamin PG. The chemokine receptor CX3CR1 mediates homing of MHC class II-positive cells to the normal mouse corneal epithelium. Invest Ophthalmol Vis Sci 2007;48:1568–1574.

66. Ishida Y, Gao JL, Murphy PM. Chemokine receptor CX3CR1 mediates skin wound healing by promoting macrophage and fibroblast accumulation and function. J Immunol 2008;180:569–579.

67. Pham TL, Kakazu A, He J, Bazan HEP. Mouse strains and sexual divergence in corneal innervation and nerve regeneration. FASEB J 2019;33:4598–4609.

68. Kwon B, Hazlett LD. Association of CD4+ T cell-dependent keratitis with genetic susceptibility to Pseudomonas aeruginosa ocular infection. J Immunol 1997;159:6283–6290.

69. Hazlett LD, McClellan S, Kwon B, Barrett R. Increased severity of Pseudomonas aeruginosa corneal infection in strains of mice designated as Th1 versus Th2 responsive. Invest Ophthalmol Vis Sci 2000;41:805–810.

70. Berk RS, Leon MA, Hazlett LD. Genetic control of the murine corneal response to Pseudomonas aeruginosa. Infection and immunity 1979;26:1221–1223.

71. Berk RS, Beisel K, Hazlett LD. Genetic studies of the murine corneal response to Pseudomonas aeruginosa. Infection and immunity 1981;34:1–5.

72. Li F, Yu R, Sun X, et al. Autonomic nervous system receptor-mediated regulation of mast cell degranulation modulates the inflammation after corneal epithelial abrasion. Exp Eye Res 2022;219:109065.

73. Xue Y, He J, Xiao C, et al. The mouse autonomic nervous system modulates inflammation and epithelial renewal after corneal abrasion through the activation of distinct local macrophages. Mucosal Immunol 2018;11:1496–1511.

74. He S, Liu J, Xue Y, Fu T, Li Z. Sympathetic Nerves Coordinate Corneal Epithelial Wound Healing by Controlling the Mobilization of Ly6Chi Monocytes From the Spleen to the Injured Cornea. Invest Ophthalmol Vis Sci 2023;64:13.

75. Xiao C, Wu M, Liu J, et al. Acute tobacco smoke exposure exacerbates the inflammatory response to corneal wounds in mice via the sympathetic nervous system. Commun Biol 2019;2:33.

76. Yun H, Yee MB, Lathrop KL, Kinchington PR, Hendricks RL, St Leger AJ. Production of the Cytokine VEGF-A by CD4(+) T and Myeloid Cells Disrupts the Corneal Nerve Landscape and Promotes Herpes Stromal Keratitis. Immunity 2020;53:1050–1062 e1055.

77. Ma X, Wang Q, Song F, et al. Corneal epithelial injury-induced norepinephrine promotes Pseudomonas aeruginosa keratitis. Exp Eye Res 2020;195:108048.

78. Li J, Ma X, Zhao L, Li Y, Zhou Q, Du X. Extended Contact Lens Wear Promotes Corneal Norepinephrine Secretion and Pseudomonas aeruginosa Infection in Mice. Invest Ophthalmol Vis Sci 2020;61:17.

79. Zhang BN, Qi B, Chu WK, et al. Norepinephrine as the Intrinsic Contributor to Contact Lens-Induced Pseudomonas aeruginosa Keratitis. Invest Ophthalmol Vis Sci 2023;64:26.

80. Li W, Gurdziel K, Pitchaikannu A, Gupta N, Hazlett LD, Xu S. The miR-183/96/182 cluster is a checkpoint for resident immune cells and shapes the cellular landscape of the cornea. The ocular surface 2023;30:17–41.

